# Regulatory regions in natural transposable element insertions drive interindividual differences in response to immune challenges in Drosophila

**DOI:** 10.1101/655225

**Authors:** Anna Ullastres, Miriam Merenciano, Josefa González

## Abstract

**Background:** Variation in gene expression underlies interindividual variability in relevant traits including immune response. However, the genetic variation responsible for these gene expression changes remain largely unknown. Among the non-coding variants that could be relevant, transposable element insertions are promising candidates as they have been shown to be a rich and diverse source of cis-regulatory elements.

**Results:** In this work, we used a population genetics approach to identify transposable element insertions likely to increase the tolerance of *Drosophila melanogaster* to bacterial infection by affecting the expression of immune-related genes. We identified 12 insertions associated with allele-specific expression changes in immune-related genes. We experimentally validated three of these insertions including one likely to be acting as a silencer, one as an enhancer, and one with a dual role as enhancer and promoter. The direction in the change of gene expression associated with the presence of several of these insertions was consistent with an increased survival to infection. Indeed, for one of the insertions, we showed that this is the case by analyzing both natural populations and CRISPR/Cas9 mutants in which the insertion was deleted from its native genomic context.

**Conclusions:** We showed that transposable elements contribute to gene expression variation in response to infection in *D. melanogaster* and that this variation is likely to affect their survival capacity. Because the role of transposable elements as regulatory elements is not restricted to Drosophila, TEs are likely to play a role in immune response in other organisms as well.

## BACKGROUND

Gene expression changes occur across developmental stages and cell types, and in response to external stimuli and disease. Identifying the genetic variation underlying context-dependent differences in gene expression is thus essential to understand organism’s development and functioning (1–3). Experimental and computational efforts aimed at identifying gene regulatory elements are biased towards activating elements, namely enhancers and promoters, while silencing regulatory elements are less-studied. Moreover, recent advances suggest that the classical definition of regulatory elements needs to be updated. Activating regulatory elements may have both enhancer and promoter functions and silencers may act as enhancers in alternate cellular contexts (2, 4, 5). Another bias of the current approaches is that they are mostly focused on the analysis of single nucleotide variants (6). However, transposable elements (TEs) are also known to be a rich and diverse source of cis-regulatory elements (7–12).

TEs are repetitive DNA sequences with the ability to move along the genome (13). They can disperse promoters and enhancers that increase gene expression levels (14, 15) and they can also silence gene expression by spreading heterochromatin formation and stalling Pol II elongation (12, 16). TEs have also been documented to function as boundary elements or insulators and to contribute to non-coding regulatory sequences that modify gene expression post-transcriptionally (9, 11). In addition to providing cis-regulatory elements to individual genes, TEs also have an impact on gene regulation at a genome-wide level in different organisms such as human, mouse, and *D. melanogaster* (17–26). The genome-wide contribution of TEs to gene regulation has been studied in several relevant phenotypes such as development, response to xenobiotics, immune response, and disease (23, 27–29). While there are several studies highlighting the potential role of TE insertions in the regulation of the immune response (30–32), whether TEs are more often recruited to play a role in immune response remains speculative (33).

Innate immunity is the first barrier against infections, and many species rely solely on this response to cope with pathogens (34, 35). One of the most likely infection routes occurring in nature is oral infection, and the gut epithelium is the first barrier that bacteria encounter in the organism (36, 37). However, the gut immune response is still not completely understood, and it is likely more complex than the systemic immune response (36, 38–40). Variation in gene expression has been shown to underlay interindividual variability in immune responses in several organisms including humans (41–45). In *D. melanogaster*, gut immunocompetence variation was analyzed in 140 strains, and small but systematic differences in gene expression were found between resistant and susceptible strains to *Pseudomonas entomophila*, a natural pathogen of this species (41, 46). However, the causal mutations responsible for these expression changes remain largely unknown (45, 47, 48). The ability to carry out *in vivo* enhancer assays and CRISPR/Cas9 mediated genomic deletions makes *D. melanogaster* a prime choice to study the role of TEs in immune-related processes.

In this work, we aimed at identifying transposable element variants that could be contributing to the ability of *D. melanogaster* populations to cope with immune challenges. While the identification of regulatory variants is often based on sequence analysis, we used a population genetics approach in which we look for TE insertions likely to be evolving under positive selection. We then performed allele-specific expression analysis to identify which of the candidate TE insertions were associated with changes in gene expression. Finally, we combined multiple experimental techniques, including CRISPR/Cas9, to provide additional evidence of functionality for a subset of the identified candidate TE insertions.

## RESULTS

### Nineteen natural transposable element insertions present at high population frequencies are located nearby genes with immune-related functions

To identify polymorphic transposable elements (TEs) that are likely to be adaptive and to play a role in immune-response, we first looked for insertions present at both high population frequencies and in genomic regions with recombination rate larger than zero (see Methods). The rationale for this screening is as follows: (i) if a mutation is adaptive, that particular mutation is expected to be present at high population frequencies; and (ii) although slightly deleterious or neutral mutations could also increase in frequency in populations, this is less likely in regions with recombination rate larger than zero because the efficiency of selection in these regions is high (49). We analyzed the frequency of 831 TE insertions located in regions with recombination rates larger than zero, in four natural populations: Siavonga (Zambia), Stockholm (Sweden), Bari (Italy) and North Carolina (USA) (Additional File 1A and 1B, see Methods) (50–52). Although the majority of these 831 TE insertions are present in the reference genome, we also included in our analysis 23 *de novo* TE insertions (51) (see Methods). Overall, we identified 128 TE insertions present at >10% frequency in at least one of the four populations analyzed (Additional File 1C, see Methods).

We then surveyed the literature for any functional information available for the genes located nearby each of these 128 TEs (Additional File 1D). We found that 19 of these TEs were associated with 21 immune-related genes (Table 1). The functional evidence for the majority of these genes comes from transcriptional response to infection (12 genes), infection survival experiments (five genes), or both (two genes; Table 1). The other two genes, *TM4SF* and *ken*, are also involved in immune response. *TM4SF* is a member of the tetraspanin transmembrane proteins, which modulate immune signaling in *Drosophila* (53), and *ken* is a member of the JAK-STAT signaling pathway involved in immune response (54). To provide further evidence for the role of these genes in immune-related functions, we performed infection survival experiments with *Pseudomonas entomophila* for five genes for which survival experiments were not previously available (*CG2233*, *ken*, *CG8008*, *TM4SF* and *CG10943*), and for three genes for which survival experiments were performed using a different pathogen (*NUCB1*, *Bin1* and *cbx*). *P. entomophila* (46) is a natural *D. melanogaster* pathogen, and thus experiments with these gram-negative bacteria have the potential to identify specialized immune responses derived from antagonistic co-evolution (55). To perform the infection survival experiments, we used a mixture of RNAi knockdown lines, gene disruption lines, and overexpression lines, and when possible, we used two different genetic backgrounds (Figure 1, Additional File 2A). We first confirmed that the RNAi knockdown lines, gene disruption lines, and overexpression lines indeed showed changes in expression of the target genes (Additional File 2A and 2B; see Methods). We found that changes in expression for seven of the eight genes tested were associated with differences in survival after infection: expression changes in *NUCB1, CG2233, Bin1* and *cbx* were associated with higher survival, while expression changes in *ken*, *CG8008*, and *TM4SF* were associated with lower survival (Figure 1, Additional File 2C). For *CG10943* and for one of the two backgrounds tested for *Bin1*-although we found differences in survival-the effect size was not significant (Figure 1, Additional File 2C).

**Table 1.** Candidate TEs located nearby immune-related genes. TE length is indicated as full-length (FL) or truncated (T). In the case of truncated elements, whether the insertion is a solo LTR, a TIR or 5’ truncated (5’) is also indicated. NS: not significant. NA: not analyzed.

| TE | TE family (class) | TE start position | TE length | Evidences of selection | Gene (position) | Immune-related evidence |
| --- | --- | --- | --- | --- | --- | --- |
| *FBti0019457* | pogo (DNA) | 3R: 29760415 | 1,146 bp (T: TIR) | Fst, nSL (52) | *kay* (4,434 bp 5') | **Expression.** Component of the JNK pathway, essential for antimicrobial peptide release (118, 119). Up-regulated in *imd* and *bsk* mutant LPS-induced S2 cells, and down-regulated in *Rel* mutants (120). Up-regulated in larvae infected with *P. entomophila* (gram-negative) (46), and after 4h of infection with *P. entomophila* (41). |
| *FBti0020046* | Doc (non-LTR) | 3L: 6040416 | 2,305 bp (T: 5') | Allele age (121) | *Jon65Aiv* (281 bp 3') | **Expression.** Up-regulated after septic injury with mixed bacteria: *M. luteus* (gram-positive) and *E. coli* (gram-negative) (122). Down-regulated after 4h of infection with *P. entomophila* (41). Down-regulated after oral infection with the avirulent *gacA P. entomophila* (123). |
| *FBti0020057* | BS (non-LTR) | 3L: 7130011 | 126 bp (T: 5') | H12, nSL (52) | *CG15829* (338 bp 3') | **Expression.** Up-regulated after infection by septic injury with mixed bacteria (gram-positive and gram-negative), and regulated by *Rel* (122). Up-regulated after 4h of infection with *P. entomophila* (41). Up-regulated in transgenic flies expressing a SINV replicon (alphavirus) (124). |
|  |  |  |  |  | *CG8628* (739 bp 5') | **Expression.** Up-regulated in microbiota associated flies vs germ free flies (125), after infection with several pathogens (gram-positive and gram-negative, fungi, protozoa) (126), and down-regulated after 4h of infection with *P. entomophila* (41). |
| *FBti0018883* | Burdock (LTR) | 2R: 9151357 | 6,413 bp (T: LTR) | NS | *CG8008* (136 bp 3') | **Expression.** Induced by LPS (gram-negative) in an IKK-dependent manner in S2 cell cultures (127). Up-regulated after *E. coli* (gram-negative) infection in S2 cells (128). |
| *FBti0019381* | Juan (non-LTR) | 3R:<br>15132112 | 2,995 bp<br>(T: 5') | NS | *CG42788*<br>(180 bp<br>5') | **Expression.** Down-regulated in response to *P. rettgeri* (gram-negative) infection in females (129). |
| *FBti0019602* | Juan (non-LTR) | X: 8031495 | 4,249 bp<br>(FL) | NS | *CG2233*<br>(12 bp 3') | **Expression.** Down-regulated in *PEBP1* mutant L3 larvae, which are more resistant to *M. luteus* (gram-positive) and *E. coli* (gram-negative) infection (130). Latitudinal expression differentiation after infection with *E. coli* and *M. luteus* mix in temperate vs. tropical populations (65). |
| *FBti0061105* | G5 (non-LTR) | 2R: 7317828 | 51 bp<br>(T: 5') | NS | *Dscam1*<br>(46 bp 3') | **Expression.** Required in hemocytes for efficient phagocytosis and binds to *E. coli* (gram-negative) (131). |
| *FBti0062242* | BS (non-LTR) | 3R:<br>16041234 | 102 bp<br>(T: 5') | NS | *Pnr*<br>(3'UTR) | **Expression.** *pnr* is a modifier of the Toll pathway and RNAi mutants show Imd pathway hyperactivation when infected with *E. cloacae* (gram-negative) and *M. luteus* and *E. faecalis* (gram-positive) (132). |
| *tdn4* | BS (non-LTR) | 2R:<br>18807871 | 800 bp<br>(T: 5') | NA | *CG15096*<br>(479 bp<br>3') | **Expression.** Down-regulated in Oregon R and *Rel*-mutant flies with microbiota compared to axenic flies (133), and after *P. entomophila* infection (41). |
| *tdn8* | Transpac (LTR) | 3L:<br>12863675 | 5,500 bp<br>(FL) | NA | *CG10943*<br>(816 bp<br>5') | **Expression.** Up-regulated in Oregon R and *Rel*-mutant flies with microbiota compared to axenic flies (133), 24h after infection with *O. muscaedomesticae* (protozoan) (126), and after *P. entomophila* infection (41). |
| *tdn17* | pogo (DNA) | X:<br>21399382 | 2,067 bp<br>(FL) | NA | *lcs* (2,067<br>bp 5') | **Expression.** Involved in virus response, down-regulated in males infected with sigma virus (134). Up-regulated in young fly guts (133). |
| *FBti0019386* | invader4 (LTR) | 3R:<br>16189464 | 347 bp<br>(T: solo<br>LTR) | CL test,<br>TajimaD,<br>Phenotypic<br>(135) | *Bin1*<br>(5'UTR) | **Survival.** Mutant larvae are more sensitive to fungal *A. fumigatus* (fungi) infection (136). |
| *FBti0019985* | roo (LTR) | 2R: 9871090 | 434 bp<br>(T: solo<br>LTR) | TajimaD,<br>iHS, H12,<br>Phenotypic<br>(137) | *cbx* (first<br>intron) | **Survival.** Mutant flies are more sensitive to *S. aureus* (gram-positive) septic infection, but not to *S. typhimurium* (gram-negative) infection (58). |
| *FBti0019564* | mdg1 (LTR) | X: 3785867 | 189 bp<br>(T: LTR) | TajimaD<br>(138) | *tlk*<br>(intron) | **Survival.** Involved in antimicrobial humoral response to gram-negative (119). *tlk* knockdown, together with other five genes knocked-down, reduces phagocytosis of *E. coli* (gram-negative) and *S. aureus* (gram-positive) in S2 cells (139). |
| *FBti0020119* | S (DNA) | 3L:<br>15554974 | 1,732 bp<br>(FL) | NS | *AGO2*<br>(first<br>intron) | **Survival.** Involved in defense response to virus infections (140, 141), and interacts with Imd pathway proteins during gram-negative infection (142). |
| *FBti0020137* | S (DNA) | 3L:<br>17799864 | 1,732 bp<br>(FL) | NS | *NUCB1*<br>(first<br>intron) | **Survival.** Mutants are more resistant to *V. cholerae* (gram-negative) oral infection (59). |
| *FBti0061506* | 1360 (DNA) | 2L:<br>17432071 | 48 bp<br>(T: TIR) | iHS (52) | *Dif* (first<br>intron) | **Survival and expression.** Transcription factor involved in defense response to fungus and gram-positive bacteria and mediates Toll pathway activation (143-147). *Dif* mutants are susceptible to fungi and gram-positive bacterial infection (147, 148). Up-regulated in guts from *P. entomophila* infected flies (41). |
| *FBti0018877* | BS (non-LTR) | 2R: 9945496 | 131 bp<br>(T: 5') | NS | *Mef2*<br>(first<br>intron) | **Survival and expression.** Adult *Mef2* mutant males are more sensitive to *E. cloacae* (gram-negative) and *M. marinum* (gram-positive) septic infection (149). Up-regulated after 4h of infection with *P. entomophila* (gram-negative) (41). |
| *FBti0018868* | 297 (LTR) | 2R:<br>23877783 | 414 bp<br>(T: LTR) | NS | *TM4SF*<br>(1 bp 5') | A **tetraspanin** transmembrane protein, which modulate immune-signaling in *Drosophila* (53). |
|  |  |  |  |  | *ken* (340<br>bp 3') | **JAK-STAT.** Member of JAK-STAT pathway (150). JAK-STAT pathway plays a role in immune response in *D. melanogaster* (54). |

**Figure 1.**
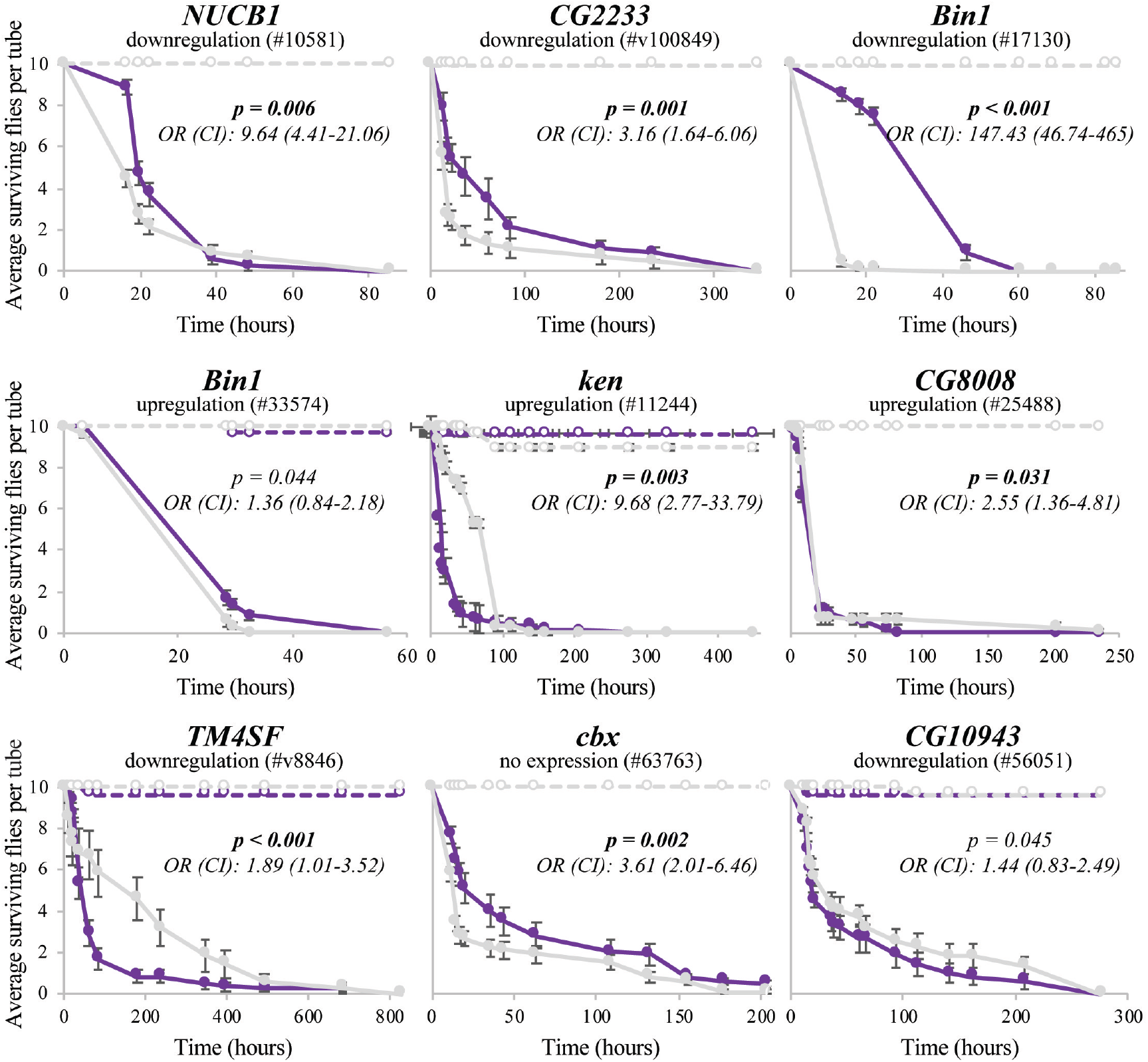
*NUCB1*, *CG2233*, *Bin1*, *ken*, *CG8008*, *TM4SF*, and *cbx* are associated with changes in survival rates after an oral infection with *P. entomophila*. Survival curves in non-infected (discontinuous lines) and infected conditions (continuous lines) for gene disruption, RNAi, and overexpressing flies for each gene (purple) and wild-type flies (grey). Error bars represent the standard error of the mean (SEM). Log-rank p-values, odds-ratios (OR), and 95% confidence intervals (CI) are given for each survival assay.

Overall, we identified 19 candidate adaptive TEs located near 21 immune-related genes. While for some of these genes there is previous evidence linking changes in expression with changes in survival to infection (Table 1), we provided additional evidence for seven genes further suggesting that they do play a role in immune response (Figure 1, Additional File 2B and 2C). Note that there is prior evidence suggesting that seven of these 19 TEs have increased in frequency due to positive selection (Table 1) (52). We next investigated whether the identified candidate adaptive TEs were associated with changes of expression of their nearby immune-related genes.

### Immune-related candidate TEs are associated with gene expression changes

In order to explore whether the 19 candidate adaptive TEs were associated with expression changes of their nearby immune-related genes, we measured allele-specific expression (ASE) in flies heterozygous for the presence of each candidate adaptive TE. Because both alleles in the heterozygote share the same cellular environment, differential expression of the two alleles are indicative of functional cis-regulatory differences (56, 57). We performed the analysis in flies with two different genetic backgrounds in order to detect possible background-dependent effects in allele-specific expression changes.

We were able to analyze a total of 16 genes located nearby 14 TEs (see Methods). In non-infected conditions, 10 out of the 16 genes showed statistically significant allele-specific expression differences in at least one of the two genetic backgrounds analyzed (Figure 2, Additional File 3). For five of these genes, we found that the allele with the TE was more highly expressed compared with the allele without the TE, and for the other five genes, the allele with the TE was less expressed. In infected conditions, eight out of the 16 genes showed statistically significant allele-specific expression differences in at least one of the two genetic backgrounds analyzed (Figure 2, Additional File 3). For three of these genes, we found that the allele with the TE was more highly expressed, and for the other five genes, the allele with the TE was less expressed. Note that up-regulation of *CG10943* and *Dif* and down-regulation of *CG8628, CG15096, NUCB1* and *cbx* in infected conditions has been previously associated with increased tolerance to infection (Table 1) (41, 58, 59).

**Figure 2.**
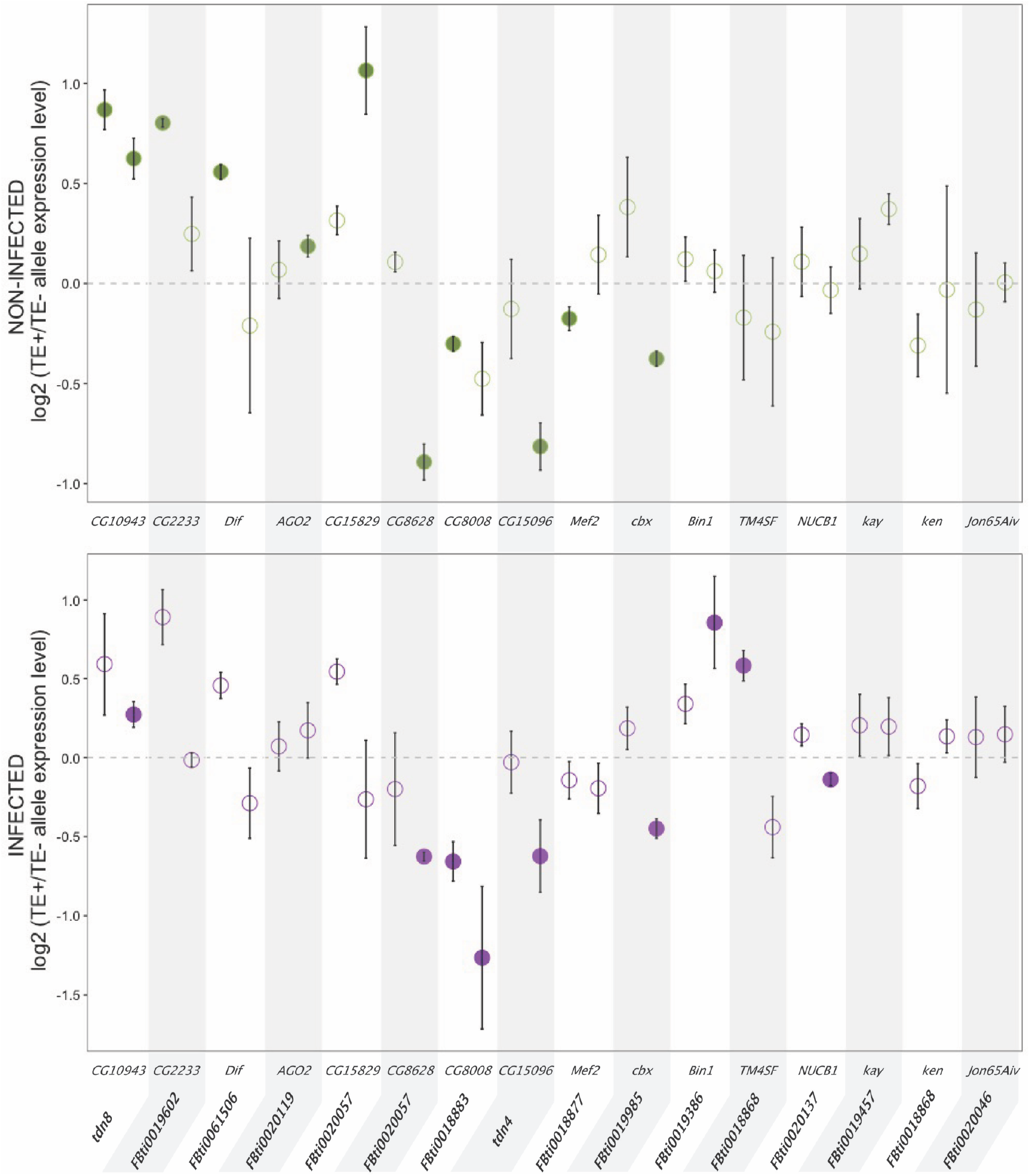
Twelve candidate immune-related TEs are associated with changes in expression of their nearby genes. Allele specific expression results from female guts in non-infected conditions (top, green) and in infected conditions (bottom, purple). Each dot represents the average ratio of gene expression levels between the allele with the TE and the allele without the TE for the three replicates analyzed. Each gene has two dots representing each one of the two genetic backgrounds analyzed. Statistically significant differences are depicted with darker colors (t-test p-values < 0.05, corrected for 5% FDR). Error bars represent SEM. Note that besides being associated with lower *CG8628* allele-specific expression, *FBti0020057* was also associated with increased *CG15829* allele-specific expression in non-infected conditions.

Considering both non-infected and infected conditions, five genes showed allele-specific expression differences under both conditions: for *CG10943* the allele with the TE was more highly expressed, and for *CG8628*, *CG8008*, *CG15096* and *cbx* the allele with the TE was less expressed (Figure 2, Additional File 3).

We also checked whether the genetic background affected the allele specific expression differences. In ten of these comparisons, both backgrounds showed changes in expression in the same direction, and in two of them the differences were statistically significant in the two backgrounds analyzed (Figure 2, Additional File 3). On the other hand, while seven analyses showed differential expression in opposite directions in the two backgrounds, results were always only statistically significant in one of the two backgrounds analyzed (Figure 2, Additional File 3).

Finally, we checked whether there were polymorphisms linked to the presence of the TE insertion that could also be associated with the detected differences in allele expression (see Methods). Only for the *AGO2* gene, we found two single nucleotide polymorphisms (SNPs) in the coding region that were linked to *FBti0020119* insertion (Additional File 6B). Thus, and although we cannot completely exclude that other polymorphisms could also be playing a role, the TE insertion appeared to be the most likely causal mutation with the exception of *FBti0020119*.

Overall, we found that most of the candidate immune-related TEs, 12 out of 14, were associated with changes in expression of their nearby gene, in at least one of the two conditions analyzed (Figure 2). While some expression changes are significant only in infected or only in non-infected conditions, a significant proportion of genes (38%) showed consistent changes in expression in both conditions (Figure 2). We also detected an effect of the genetic background on the allele specific expression differences as has been previously reported (60–62). Finally, while recent studies performed in several *D. melanogaster* strains estimated that ~8% to 28% of genes showed allele-specific expression in control conditions (63–65), we found this percentage to be 62.5%, which is significantly higher than the previous reported higher value (Fisher test: p-value = 0.0087).

### Candidate adaptive TEs associated with lower allele-specific expression are not enriched for the repressive histone mark H3K9me3

Six of the TEs analyzed were associated with lower allele-specific expression levels (*FBti0020057, FBti0018883, tdn4, FBti0018877, FBti0019985*, and *FBti0020137;* Figure 2). One of the molecular mechanisms by which TEs are associated with gene down-regulation is by recruiting repressive histone marks such as H3K9me3 (66–68). Thus, we checked whether flies homozygous for each one of these insertions showed H3K9me3 enrichment in the region where the TE is inserted compared with strains without the insertions. We found that the strains with the insertion were not enriched in H3K9me3 histone mark compared with the strains without the insertion (p-values > 0.05) (Figure 3A). Indeed, for *FBti0019985* we found a depletion of H3K9me3 in the TE region (Figure 3A). Thus, the lower allele-specific expression associated with the TEs observed in the ASE analysis is not due to changes in H3K9me3 repressive histone mark (Figure 3A).

**Figure 3.**
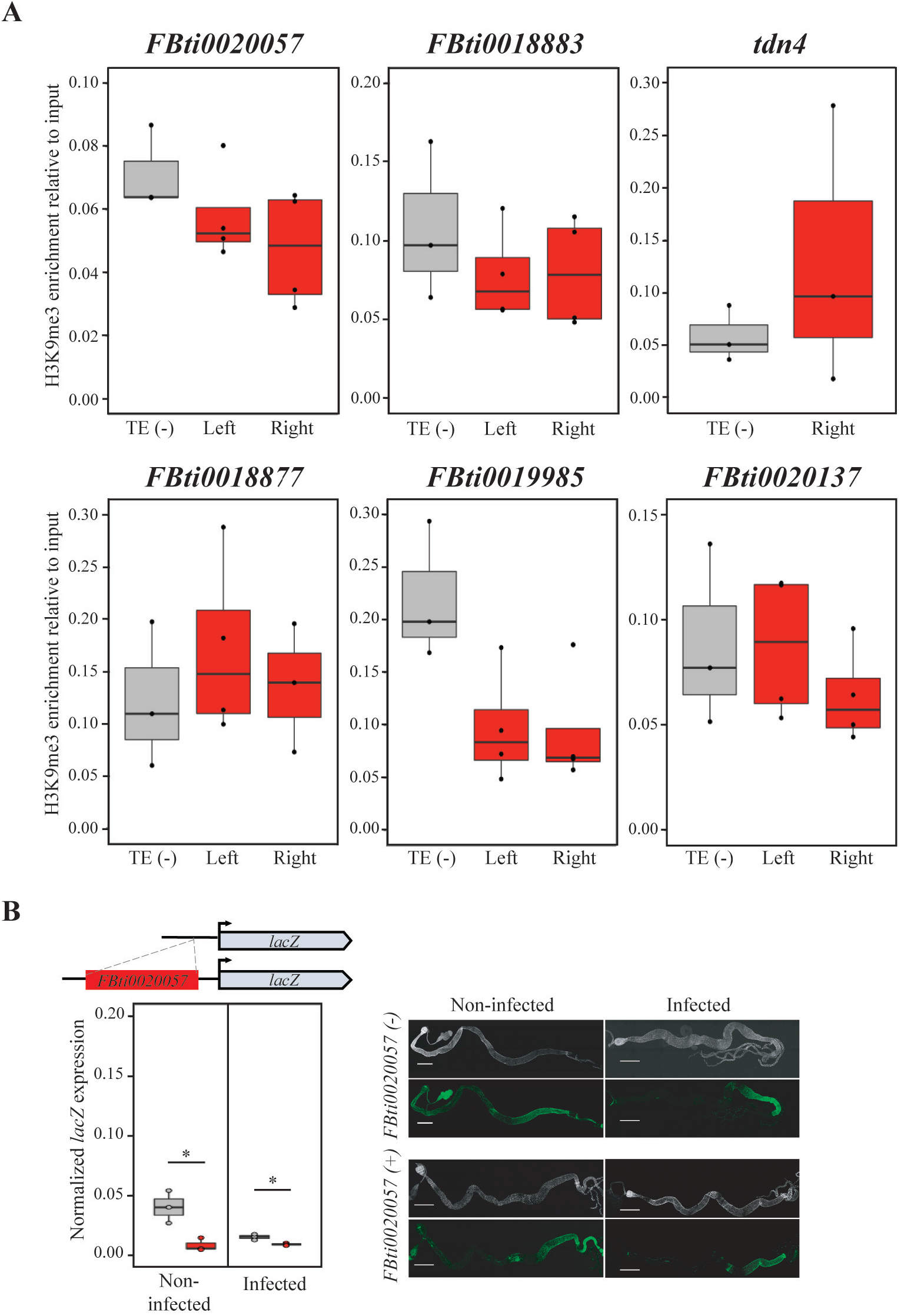
Candidate immune-related TEs associated with lower allele-specific expression are not enriched for H3K9me3. **A)** ChIP RT-qPCR analysis for H3K9me3 in the genomic region where *FBti0020057, FBti0018883, tdn4, FBti0018877, FBti0019985,* and *FBti0020137* are inserted. TE (−): H3K9me3 enrichment in the strain that does not contain the TE insertion (grey). Left: H3K9me3 enrichment in the left flanking region of each TE in a strain with the insertion (red). Right: H3K9me3 enrichment in the right flanking region of each TE in a strain with the insertion (red). None of the candidate immune-related TEs tested are enriched for H3K9me3. *FBti0020057*: p = 0.384 and p = 0.115 for the left and right TE-flanking regions, respectively; *FBti0018883*: p = 0.473 and p = 0.425, respectively; *tdn4*: p = 0.408 for the right TE-flanking region; *FBti0018877*: p = 0.364 and p = 0.632, respectively; *FBti0019985*: p = 0.041 and p = 0.039, respectively; and *FBti0020137*: p = 0.880, and p = 0.423, respectively. **B)***FBti0020057* is associated with lower reporter gene expression in both non-infected and infected conditions. Schematic representation of the vector construction with the intergenic region between *CG15829* and *CG8628* genes with and without *FBti0020057* insertion cloned upstream of the reporter gene *lacZ*. Below, normalized expression levels of the *lacZ* reporter gene in transgenic female guts with (red) and without (grey) *FBti0020057* are shown. On the right side, β-GAL immunostaining (green), and DAPI staining (grey) of guts from transgenic females with and without *FBti0020057*. Scale bars represents 500 μm.

An alternative hypothesis is that these TE insertions could be recruiting transcriptional repressors or that they are disrupting regulatory regions. To provide evidence that these TE insertions could indeed be the mutations associated with the lower allele-specific expression levels, we cloned the gene promoter region with and without a TE insertion in front of a reporter gene, using *FBti0020057* as an example. We indeed found that flies with *FBti0020057* TE insertion were associated with lower reporter gene expression both in control and infected conditions (Figure 3B; t-test, p-value = 0.019 and 0.016, respectively). Although we cannot discard that *FBti0020057* is associated with gene downregulation because it disrupts the promote gene region, it has been reported that this insertion contains a binding site for *Nub*, a transcription factor which negatively regulated many genes involved in immune and stress responses (Additional File 1E) (69).

### Two of the four tested candidate adaptive TEs associated with high allele-specific expression drives reporter gene expression under stress conditions

There is previous evidence showing that *FBti0019386*, which is associated with increased allele-specific expression in infected conditions, drives the expression of a reporter gene after oral infection with *P. entomophila* (25). To test whether other TEs associated with increased expression of their nearby genes could also be acting as enhancers, we performed *in vivo* enhancer assays. We focused on three insertions located in promoter regions (less than 1 kb from a gene) and associated with ≥ 1.5-fold increased expression: *tdn8*, *FBti0018868*, and *FBti0061506*.

We found that transgenic flies in which we cloned the upstream gene region containing the *tdn8* sequence showed increased reporter gene expression compared with transgenic flies containing the same region without the insertion (Figure 4A). Differences in gene expression were only statistically significant in infected conditions (t-test, p-value = 0.095 and 0.046 for control and infected, respectively; effect size 1.4 and 1.5 for control and infected, respectively). No differences between the two transgenic strains in the localization of the *β-GAL* protein expression in control or infected conditions were found (Figure 4A). On the other hand, we found that neither *FBti0018868* nor *FBti0061506* drive the expression of a reporter gene in non-infected or infected conditions (Figure 4B and 4C; t-test, all p-values > 0.05).

**Figure 4.**
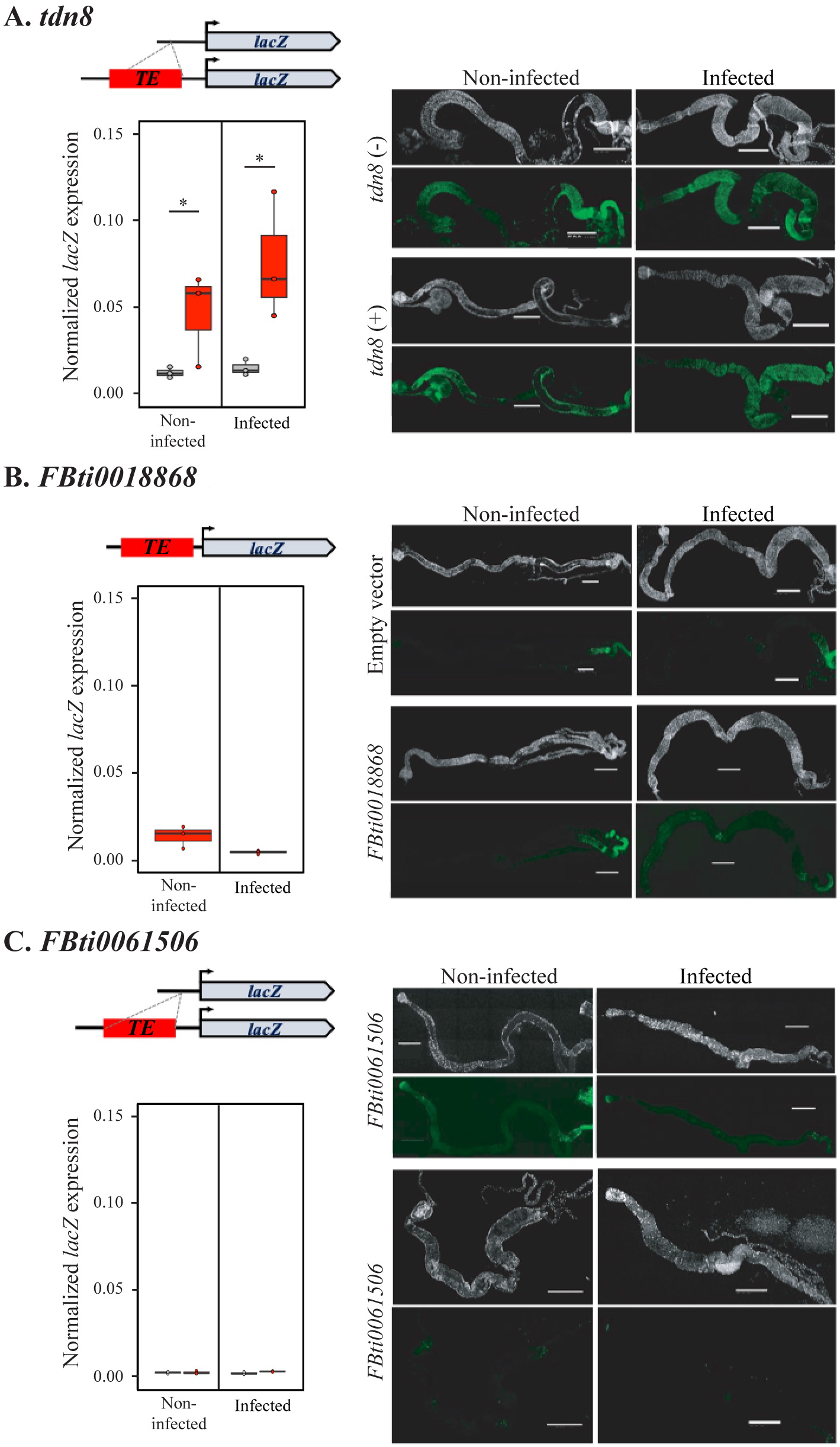
*tdn8* insertion showed increased reporter gene expression. Results of the in vivo enhancer reporter assays for **A)** *tdn8*, **B)***FBti0018868*, and **C)** *FBti0061506*. Normalized expression levels of the *lacZ* reporter gene in transgenic female guts with (red) and without (grey) the candidate TE insertions are shown. β-GAL immunostaining (green), and DAPI staining (grey) of guts from transgenic females with and without each candidate TE insertions is also shown. Scale bars represents 500 μm.

Overall, we found that two of the four tested TE insertions that are associated with ≥ 1.5-fold higher allele-specific expression drive the expression of a reporter gene (Figure 4). For the rest of this work, we focus on the *FBti0019386* insertion as a case study to further understand the molecular mechanisms and the phenotypic consequences of a candidate insertion related to immune-response.

### *FBti0019386* adds immune-related transcription factor binding sites

Villanueva-Cañas et al (2019) (25) reported that *FBti0019386*, which drives the expression of a reporter gene only in infected conditions, harbors two immune-related binding sites for *Caudal* and *DEAF-1* transcription factors (Additional File 1E) (70–72). To test whether the predicted immune-related transcription factor binding sites (TFBSs) were responsible for the enhancer activity of this insertion in infected conditions, we generated a transgenic fly line in which we deleted the two binding sites from the *FBti0019386* insertion sequence (Figure 5A). We performed expression analysis with the two previously generated transgenic lines described in Villanueva-Cañas et al (2019) (25), one containing the *Bin1* upstream region without the insertion, and one containing the *Bin1* upstream region with the insertion, and with the newly generated transgenic line in which the TFBSs were deleted.

**Figure 5.**
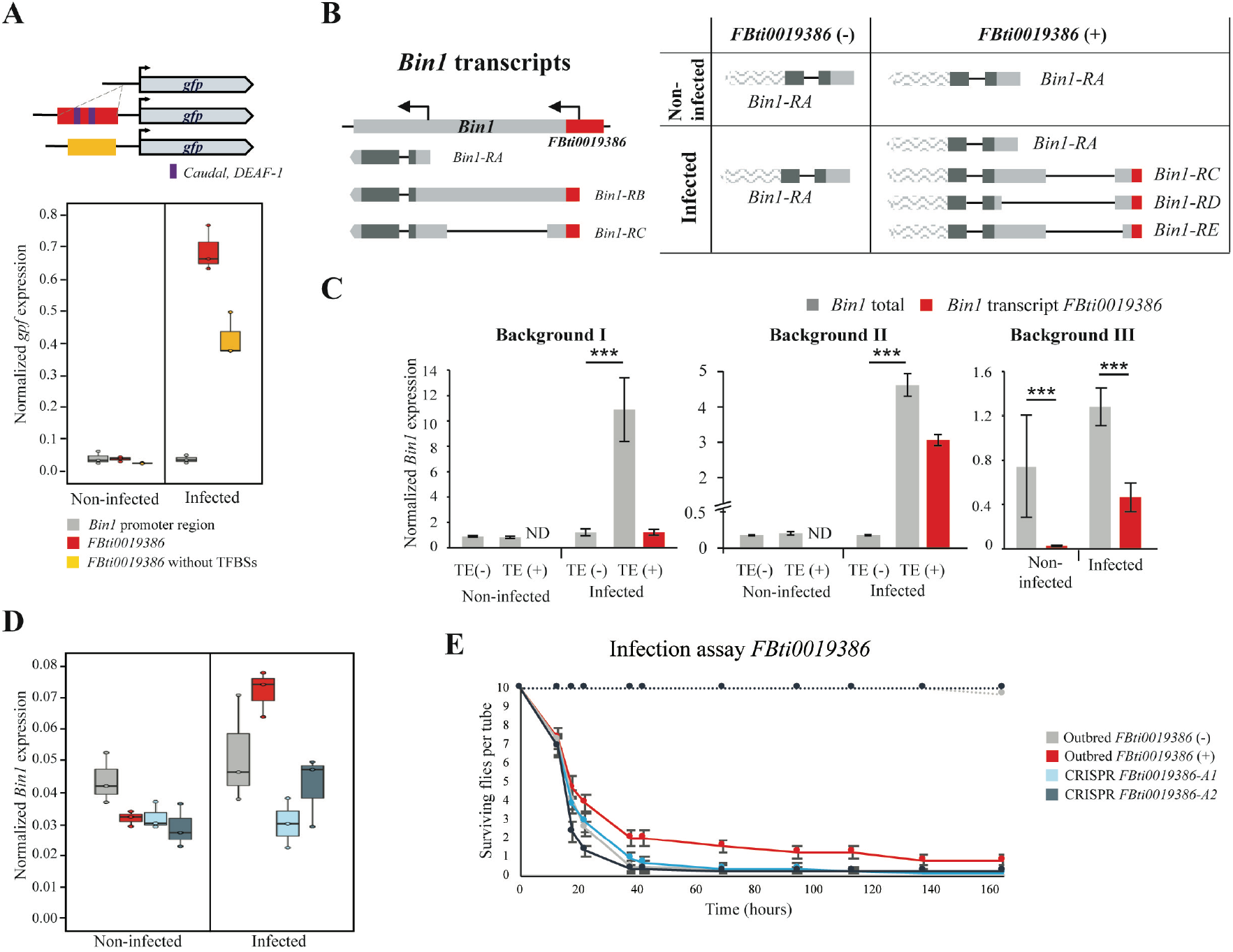
*FBti0019386* is most likely the causal mutation for the differences in expression of *Bin1* and for the increased tolerance to *P. entomophila*. **A**) *FBti0019386* harbors two immune-related TFBSs, *Caudal* and *DEAF-1*, that are responsible for the enhancer activity of this insertion in infected conditions. Schematic representation of the vector construction with the promoter region of *Bin1* without *FBti0019386* sequence, with *FBti0019386* (represented in red), and with the *FBti0019386* sequence without the immune-related binding sites for *Caudal* and *DEAF-1* (represented in yellow) cloned upstream of the reporter gene *Gfp*. Below, normalized expression levels of the reporter gene in transgenic female guts with the different constructions are shown. **B)** *FBti0019386* adds a new TSS to its nearby gene *Bin1* used under infected conditions. On the left, we represented all the transcripts annotated for *Bin1.* On the right we show the different *Bin1* transcripts found in non-infected and infected conditions in flies with and without *FBti0019386*. UTR regions are depicted in light grey while coding regions are depicted in dark grey. TE-overlapping regions are represented in red. Transcript regions wave-patterned are inferred from FlyBase transcript annotation and were not sequenced in this work. *Bin1-RD* and *Bin1-RE* transcripts are, respectively, 318 bp and 172 bp shorter compared to *Bin1-RC* transcript. **C)** Normalized *Bin1* expression levels from female guts in the two backgrounds analyzed in the ASE experiments, and in a third homozygous background with *FBti0019386* insertion (IT_Cas 11_49-5, an isofemale strain) in both non-infected and infected conditions. Error bars represent SEM. ND: not detected. **D)** Normalized *Bin1* expression levels from female guts in outbred populations without (grey) and with *FBti0019386* insertion (red), and in the two CRISPR-mutants *FBti0019386-A1*(light blue) and *FBti0019386-A2* (dark blue) in both non-infected and infected conditions. **E)** Survival curves in non-infected (discontinuous lines) and infected conditions (continuous lines) for outbred flies with *FBti0019386* insertion (red) and without this insertion (grey), and in the two CRISPR-mutants *FBti0019386-A1*(light blue) and *FBti0019386-A2* (dark blue). Error bars represent SEM.

As expected based on previous experiments, we found that guts from transgenic flies containing the TE insertion showed increased reporter gene expression compared with transgenic flies without the insertion in infected conditions (Figure 5A; t-test, p-value < 0.001) (25). Moreover, as expected if the binding sites for *Caudal* and *DEAF-1* are responsible for the increase expression of the reporter gene, the transgenic flies in which these binding sites were deleted showed decreased reporter gene expression in infected conditions (Figure 5A; t-test, p-value = 0.009).

### *FBti0019386* provides a transcription start site to*Bin1* that is only used in infected conditions in the female gut

There is also evidence showing that *FBti0019386* adds a transcription start site to its nearby gene *Bin1* (Figure 5B) (24, 73). We thus further investigated the expression levels of *Bin1* transcripts in strains with and without *FBti0019386* insertion, in control and in infected conditions. We found that female guts from homozygous flies with and without *FBti0019386* expressed only the short *Bin1-RA* transcript in non-infected conditions (Figure 5B). However, in infected conditions, flies without *FBti0019386* insertion only expressed *Bin1-RA*, while flies with *FBti0019386* expressed *Bin1-RA*, and three additional transcripts starting in the TE: *Bin1-RC*, *Bin1-RD* and *Bin1-RE* (Figure 5B). We confirmed these results by performing the experiments in a second genetic background (see Methods). Note that the *Bin1-RD* and *Bin1-RE* transcripts have not been described previously and differ in the 5’UTR length (Figure 5B).

To test whether the transcripts starting in *FBti0019386* insertion are associated with increased expression of *Bin1*, we quantified the expression of these transcripts and the total *Bin1* expression levels (Figure 5C). As expected based on our previous ASE experiments, flies with and without *FBti0019386* did not differ in *Bin1* expression levels in non-infected conditions (t-test, p-value >0.05). In infected conditions, flies with *FBti0019386* overexpressed *Bin1* compared to flies without this insertion in the two backgrounds analyzed (t-test, p-value < 0.001). The contribution of the transcripts starting in *FBti0019386* to the total *Bin1* expression is background dependent: 11.2% in background I, and 66.3% in background II (Figure 5C). To confirm that the effect is background-specific, we analyzed a third background, and we found that the TE-transcripts contributed 36.2% to *Bin1* total expression (Figure 5C). These results suggest that besides adding a transcription start site, *FBti0019386* is also affecting the expression level of the short transcript, which is consistent with the enhancer role described for this TE (Figure 5A) (25).

### *FBti0019386* appears to be the causal mutation for the differences in expression of its nearby gene

To further test that *FBti0019386* is indeed the mutation causing the observed differences in expression, we generated two outbred populations, one containing the *FBti0019386* insertion and one without this insertion. We then deleted the insertion using the CRISPR/Cas9 homology directed repair technology and established two stocks containing this deletion (*FBti0019386-A1* and *FBti0019386-A2*; see Methods). We tested the expression of *Bin1* in the two outbred and the two CRISPR/Cas9 mutant strains. We found that outbred flies with the insertions had increased *Bin1* expression levels compared with outbred flies without the insertion in infected conditions (Two-way ANOVA; genotype effect: p=0.501, treatment effect: p=0.003, interaction genotype*treatment: p=0.025) (Figure 5D). These results are consistent with the TE acting as an enhancer only in infected conditions (Figure 5A). We also found reduced *Bin1* expression levels in the two CRISPR/Cas9 mutant strains only in infected conditions, consistent with *FBti0019386* being the causal mutation (*FBti0019386-A1*: Two-way ANOVA; genotype effect: p<0.001, treatment effect: p<0.001, interaction genotype*treatment: p<0.001, and *FBti0019386-A2*: Two-way ANOVA; genotype effect: p=0.006, treatment effect: p<0.001, interaction genotype*treatment: p=0.016) (Figure 5D).

In summary, while outbred populations with *FBti0019386* insertion showed increased *Bin1* expression levels, this expression was significantly reduced when the *FBti0019386* was deleted from its genomic context (Figure 5D).

### *FBti0019386* is associated with increased tolerance to *P. entomophila* infection

To check whether the changes in *Bin1* expression most likely caused by *FBti0019386* have an effect on the fly immune response, we measured the tolerance to oral infection of *P. entomophila* in both outbred and CRISPR/Cas9 mutant strains. We found that outbred flies with *FBti0019386* showed increased tolerance to infection compared to outbred flies without the insertion (log rank; p-value = 0.049) (Figure 5E). Although both CRISPR/Cas9 mutants have reduced tolerance compared to the strain with the TE, only *FBti0019386-A2* flies showed statistically significant differences in survival (log rank; p-value = 0.060 and p-value = 0.001, respectively) (Figure 5E).

Overall, we showed that *FBti0019386* is most likely the causal mutation for the increased tolerance to bacterial infection found in natural populations containing this TE insertion.

## DISCUSSION

In this work, we identify 12 transposable element (TE) insertions associated with allele-specific expression differences in immune-related genes (Figure 2 and Figure 6). Allele-specific expression analysis is a powerful technique to identify cis-regulatory variants (56, 57). Because the two alleles studied shared the same cellular environment, differences in expression found between them are indicative of functional cis-regulatory differences. Several works have tried to quantify the relative contribution of cis-regulatory mutations to gene expression changes in *D. melanogaster* (63–65). While these studies found that between ~8% and 28% of *D. melanogaster* genes showed allele-specific expression, we found a significantly higher proportion (62.5%, Fisher test: p-value = 0.0087). While these results might be at least partly explained by immune-related genes having an elevated rate of adaptive evolution, a global survey in Drosophila suggested that rapid adaptation of immune-related genes is the exception rather than the rule (74, 75).

**Figure 6.**
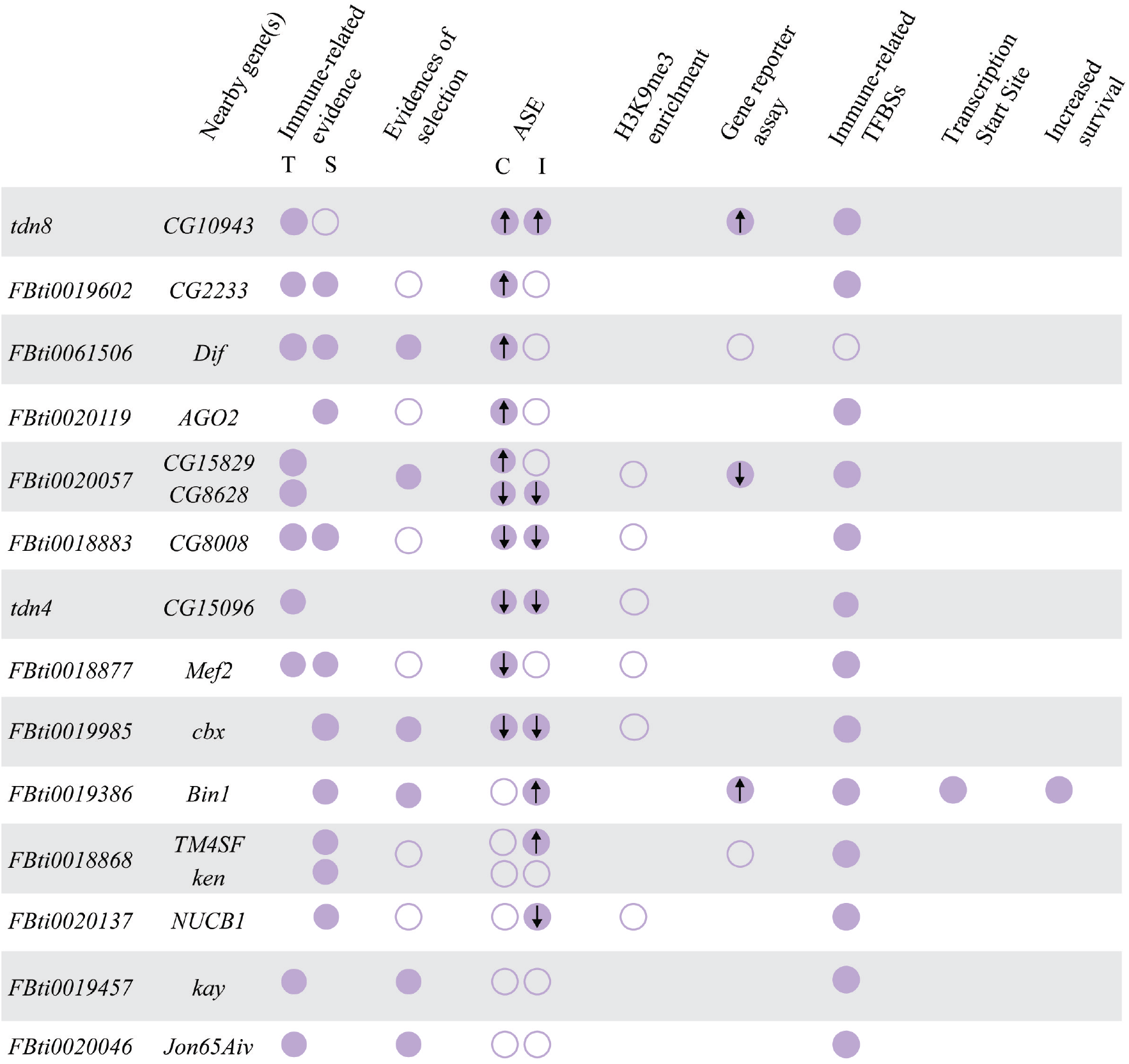
Summary of the information available for the 14 TEs associated with allele-specific expression changes. Full circles indicate statistically significant results and empty circles indicate negative or not statistically significant results. Immune-related evidence for the genes nearby includes transcriptional response to stress (T), as reported in Table 1, and survival experiments (S) from previous works (Table 1) and from experiments performed in this work (Figure 1). Evidence of selection for the regions flanking the TE insertions and immune-related transcription factor binding sites (TFBS) were obtained from the literature (25, 52, 121, 135, 137, 138). ASE: results of the allele specific expression analysis (“C” represents control conditions while “I” represents infected conditions). For ASE and for the gene reporter assays, arrows inside the circles represent the direction of the gene expression.

There is prior evidence showing that the presence of TE insertions within 1kb of a gene is associated with a larger contribution to cis-regulatory expression variation compared with genes that do not have TE insertions in their vicinity (63). However, this observation was found to be limited to genes expressed in the ovary. Our results indicate that the greater contribution of TEs to allele-specific expression also affects immune-related genes expressed in the gut. It would be interesting to test whether TEs are associated with an increased contribution to cis-regulatory changes in genes involved in other functions as well, as suggested by the enrichment of TEs nearby or inside genes involved in stress-response, behavior and developmental (52). While we focused on *P. entomophila* as an infectious agent only in female flies, it would also be interesting to use other pathogens to check whether the observed effects are pathogen-specific. Similarly, male flies may be used to know whether the effect of the candidate TEs varies, as it is known that sexual dimorphisms have an impact on pathogen-host interactions (76).

We found that some TEs were associated with gene down-regulation while others were associated with gene up-regulation (Figure 2). Similarly, no significant directional changes in expression were found in genes located nearby TEs in another study also performed in *D. melanogaster*, and in a study conducted in humans (19, 63). To provide further evidence for the regulatory role of the candidate TEs, we attempted to pinpoint the molecular mechanisms behind these expression changes. TE-induced gene silencing has been associated with epigenetic changes (66, 67, 77). Specifically, enrichment of the repressive chromatin mark H3K9me3 in the sequences flanking euchromatic TEs was associated with reduced expression of adjacent genes (66, 78). Contrary to our expectation, we did not find an enrichment of H3K9me3 in the regions flanking the six TE insertions analyzed (Figure 6). Interestingly, recent genome-wide analyses of silencer elements also failed to identify particular histone modifications associated with these regulatory sequences (4, 5, 79). Jayavelu et al (2020) found that on average only 3.3% of silencer elements overlap with H3K9me3 regions (5). Thus, the lack of histone marks does not necessarily preclude the role of these TE insertions in gene silencing. Indeed, we found that one of these insertions was associated with the down-regulation of a reporter gene (Figure 3B). *In vivo* enhancer assays also confirmed that two of the four insertions associated with gene up-regulation act as enhancers (Figure 4 and Figure 6) (25). Because enhancer reporter assays select for compact regulatory elements that can function in an autonomous manner, we cannot discard the causal role of the insertions that failed to drive reporter gene expression (80). Moreover, because enhancer assays could also lead to false positive results, they should be combined with assays in the native genomic context of the regulatory element such as for example creating a CRISPR/Cas9 mediated genomic deletion of the region under study (81). We did so for one of the TEs found to be acting as an enhancer, *FBti0019386*, and we were able to confirm that the TE is indeed responsible for the increased expression of its nearby gene (Figure 5). Furthermore, we showed that the increased expression is due to the presence of binding sites for immune-related transcription factors and to the expression of a transcript that starts in the TE insertion (Figure 5). This result is consistent with the dual role of regulatory regions as enhancers and promoters as has been previously suggested (2, 82). Note that the overall success validation rate in our *in vivo* enhancer assays is similar to that of other studies not focused on TE insertions (60%) (83, 84).

Finally, we also investigated whether the TE-induced changes in these genes could be relevant for the fly ability to cope with infections. Both gene up- and down-regulation have been previously related to *D. melanogaster* gut immune response (38, 46). Similarly, changes in basal gene expression, as we found in this work, also affect the susceptibility of the flies to immune-challenges (41, 85). If we focus on particular genes, up-regulation of *CG10943* and *Dif* and down-regulation of *CG8628, CG15096, NUCB1* and *cbx* in infected conditions has been associated with increased tolerance to infection (41, 58, 59). Because we found that *tdn8*, *FBti0061506*, *FBti0020057*, *tdn4*, *FBti0020137*, and *FBti0019985* induced expression changes in these genes in the same direction, we argue that it is likely that these TEs will also be associated with increased tolerance to infection (Figure 2). Indeed, changes in the expression of all the genes located nearby the 12 TEs identified in this work have been associated with differences in survival to infection (Figure 1, Table 1). For one of them, *FBti0019386*, we performed survival experiments with natural populations and with CRISPR/Cas9 mutant strains in which the TE was deleted, and we found that this insertion is associated with increased survival (Figure 5).

## CONCLUSIONS

We showed that TEs contribute to gene expression variation during infection in *D. melanogaster* and that this variation is likely to affect the fly infection survival capacity. Because the role of TEs as regulatory elements is not restricted to Drosophila, TEs are likely to be key players in immune response in other organisms as well (14, 86). For example, in humans, over 80% of genetic variants associated with infection risk map to the non-coding genome (87). Among the structural variants likely to play a role, TE insertions are good candidates as they have been found to be associated with expression changes in immune-related genes in human populations (88).

## METHODS

### Fly strains

#### DGRP strains

141 DGRP strains (89) were used to estimate the frequencies of TEs annotated in the *D. melanogaster* reference genome (see below) (Additional File 4). Besides, we used 37 DGRP strains to analyze by PCR a subset of TEs not annotated in the reference genome (51). Finally, DGRP strains were also used to perform allele specific expression analyses (ASE), detection of alternative transcripts, and enhancer assays (Additional File 4). Note that it has previously been shown that differences in the presence/absence of the endosymbiont *Wolbachia*, differences in commensal bacteria and/or feeding behavior has no major effect in the susceptibility of DGRP strains to *P. entomophila* infection (41).

#### African strains

Frequency estimates for reference TE insertions for a subset of 66 African strains collected in Siavonga (Zambia) (90) with no evidence of cosmopolitan admixtures were obtained from Rech et al (2019) (52) (Additional File 4).

#### European strains

Frequency estimates for reference TE insertions for 73 European strains, 57 from Stockholm (Sweden) and 16 from Bari (Italy), were obtained from Rech et al (2019) (52) (Additional File 4). Additionally, one strain from Bari (IT_Cas 11_49-5) was used for ASE and detection of alternative transcripts experiments and one strain from Munich (DE_Mun_15_8) was used for ASE experiments (Additional File 4).

#### Gene disruption, RNAi knockdown, and overexpression strains

We used two RNAi strains from the VDRC stock center (Additional File 2A). To generate the mutants, we crossed the strains carrying the RNAi controlled by an UAS promoter with flies carrying a GAL4 driver (a transcription activation system) to silence genes ubiquitously. We performed the experiments with F1 flies that were obtained from each cross. Based on the phenotypic markers, we separated the RNAi flies from the rest of the F1 that do not carry the GAL4 driver. The flies without the GAL4 driver were used as the baseline of the experiment. We also used nine mutant strains generated with different transposable element insertions and two overexpression strains. In this case, we used strains with similar genetic backgrounds as the baseline of the experiments (Additional File 2A).

#### Outbred strains

We generated present and absent outbred strains for *FBti0019386*. First, we selected all the strains that were present or absent for this TEs based on data generated by *T-lex2* in the DGRP, Zambia, Sweden, and Italy populations (52). Then, in these selected strains, we checked by PCR the presence/absence of other nine TEs identified is this work as they are likely to be involved in the immune response (*FBti0019985*, *FBti0061506*, *FBti0019602*, *FBti0020119*, *FBti0018883*, *FBti0018877*, *FBti0020137*, *tdn4*, and *tdn8*). For generating both TE-present and TE-absent outbred populations we selected six strains with the insertion and six strains without the insertion, respectively (Additional File 4). Moreover, present and absent outbred populations have similar frequencies of all the other 10 TEs likely to be involved in immune responses in order to not mask the effect of the studied TE. For every outbred population, we placed 10 males and 10 virgin females of each selected strain in a cage with fresh food. We maintained the population by random mating with a large population size for over four generations before starting the experiments.

### CRISPR/Cas9 mutant strains

Guide RNAs (gRNAs) were designed in the *FBti0019386* flanking region using the Fly CRISPR Target Finder (http://targetfinder.flycrispr.neuro.brown.edu) and cloned into pCFD5 plasmid following the pCFD5 cloning protocol (www.crisprflydesign.com) using the primers 5’-gcggcccgggttcgattcccggccgatgcagaagatctagaattagatatgttttagagctagaaatagcaag-3’ and 5’-attttaacttgctatttctagctctaaaaccacttcgtgattaattctgatgcaccagccgggaatcgaaccc-3’(91). To precisely delete the TE sequence, a donor DNA containing two homology arms flanking the *DsRed* sequence for homology repair were cloned into the pHD-ScarlessDsRed plasmid using Q5^®^ High-Fidelity DNA Polymerase (New England Biolabs). Left homology arm contained the sequence in 3R:16188287-16189463, while right homology arm contained the sequence in 3R:16189811-16191083 (Release 6) from the outbred population with *FBti0019386*. To avoid cleavage of the donor construct and mutagenesis after integration by CRISPR/Cas9, two single-nucleotide synonymous substitutions (G > T for sgRNA1 site; G > T for sgRNA2 site) were introduced into the two sgRNA target site PAM sequences, respectively. The pCFD5 plasmid containing the gRNAs, the donor pHD-ScarlessDsRed plasmid containing the homology arms, and a plasmid containing Cas9 endonuclease were co-injected as a unique mix into approximately 550 pre-blastoderm embryos from the outbred population with *FBti0019386*. All the injections were performed using the following mix concentrations: pCFD5 plasmid at 100ng/ul, donor plasmid at 500ng/ul, and Cas9 plasmid at 250ng/ul. Offspring was screened for eye fluorescence, and flies containing the deletion were backcrossed with the parental line for a minimum of five generations. Then, two homozygous strains containing the deletion of *FBti0019386* were established: *FBti0019386-A1* and *FBti0019386-A2*. The deletion was checked by PCR with the primers 5’-tttggaatcaatcacatcaaccc -3’ and 5’-caatgtcctgggtgtaagtctcg −3’. PCR bands were confirmed by Sanger sequencing.

### Transposable element datasets

#### TEs annotated in the reference genome

There are 5,416 TEs annotated in the release 6 of the *D. melanogaster* reference genome (73). In this work, we focused on polymorphic TEs present at high population frequencies and located in regions of the genome with recombination rate larger than zero (49). Most TE insertions are expected to be deleterious. Due to its big effective population size, we expect most TE insertions to be present at low frequencies in *D. melanogaster*. Thus, TEs present at high population frequencies are likely to be adaptive. We did not consider the 2,234 INE-1 insertions that are fixed in *D. melanogaster* populations (92–94). We also discarded 1,561 TEs that are flanked by simple repeats, nested TEs, or TEs that are part of segmental duplications because frequencies cannot be accurately estimated for these TEs using *T-lex2* (50). Finally, we discarded 813 TEs present in genomic regions with a recombination rate = 0 according to Fiston-Lavier et al (2010) (95) or Comeron et al (2012) (96). TEs present at low recombination regions are more likely to be linked to an adaptive mutation rather than being the causal mutation (97–100). Moreover, the efficiency of selection is low in these regions and, thus, slightly deleterious TEs could have reached high frequencies (101, 102). Hence, we ended up with a dataset of 808 annotated TEs for which we estimated their population frequencies using *T-lex2* (50) (Additional File 1A).

231 of the 808 annotated TEs were fixed in the four populations studied. Although some of these fixed TEs might be adaptive, we did not consider them as we cannot perform comparative functional experiments between flies with and without the insertions. We considered high frequent TEs those present at a population frequency ≥ 10%: 109 TEs. Note that varying this threshold does not substantially alter the number of TEs present at high frequencies (e.g. 95 TEs if we consider ≥ 15%).

#### Non-reference TE insertions

We also analyzed a subset of TEs identified by Rahman et al (2015) (51) in DGRP strains that are not annotated in the reference genome (Additional File 1B). We analyzed 23 TEs that are present in regions with recombination rate > 0 (95, 96), and were inferred to be present in at least 15 DGRP strains out of the 177 strains analyzed by Rahman et al (2015) (51). We obtained from Bloomington Drosophila Stock Center (BDSC) all the strains carrying each of the 23 insertions, and we confirmed by PCR the presence of the insertions in several strains (see below). For each TE, we sequenced at least one of the PCR products to confirm the presence and the family identity of the TE. For those insertions that we could verify, we estimated the frequency of each TE based on TIDAL results in the 177 DGRP strains and considered as high frequent those present at a population frequency ≥ 10%.

### Presence/Absence of TEs in the analyzed strains

We performed PCRs to confirm the *in silico* results obtained with *T-lex2* (50) and *TIDAL* (51). We designed specific primers for each analyzed TE using the online software Primer-BLAST (103) (Additional File 5). Briefly, we designed a primer pair flanking the TE (FL and R primers), which produces a PCR product with different band sizes when the TE is present and when the TE is absent. For those TEs that are present in the reference genome, we also designed a primer inside the TE sequence (L primer) that, combined with the R primer, only amplifies when the TE is present (104). To perform the PCRs, genomic DNA was extracted from 10 females from each analyzed strain.

### Functional annotation of genes nearby candidate adaptive TEs

We looked for functional information of the genes associated to the TEs present at high population frequencies using FlyBase (73). We considered all the genes that were located less than 1kb from the TEs. If the TEs did not have any gene located in the 1kb flanking regions, we considered only the closest gene. We considered GO annotations based on experimental evidence, and we also obtained functional information based on the publications cited in FlyBase. Several lines of evidence were considered: genome-wide association studies in which SNPs in the analyzed genes were linked to a phenotypic trait, differential expression analyses, and phenotypic evidence based on the analyses of mutant strains (Additional File 1D).

### *P. entomophila* infection

We infected 5- to 7- day-old female flies with the gram-negative bacteria *P. entomophila* (46). Flies were separated into food vials under CO_2_ anesthesia two days before the bacteria exposure, and were kept at 25°C. The experiments were performed as described in Neyen et al (2014) (105). Briefly, flies were starved for two hours and then they were flipped to a food vial containing a filter paper soaked with 1.25% of sucrose and bacterial pellet. The bacterial preparation was adjusted to a final OD_600_ = 100, corresponding to 6.5 ×10^10^ colony forming units per ml (106). Flies were kept at 29°C (only 12 hours for gene expression analysis) and 70% humidity, which are the optimal infection conditions for *P. entomophila*. In parallel, we exposed non-infected flies to sterile LB with 1.25% sucrose.

### Gene expression analysis

For RNA extraction, three replicates of 20-30 5 to 7 day-old females, males, or female guts from each gene disruption, RNAi, overexpressing and wild-type strain were flash-frozen in liquid nitrogen and stored at −80°C until sample processing (Additional File 2A). For each gene, RNA extraction was performed in the tissue and sex that FlyBase database reported a higher gene expression (Additional File 2A). For gene expression analysis after infection (in isofemale/inbred, outbred, and CRISPR/Cas9 mutant strains), we dissected 20-30 guts from both non-infected and infected 5 to 7 day-old females. Flies were infected with the gram-negative bacteria *P. entomophila* as mentioned above, and they were dissected after 12 hours of bacterial exposure. Samples were frozen in liquid nitrogen and stored at −80°C until sample processing. RNA was extracted using the GenElute™ Mammalian Total RNA Miniprep Kit (Merck) following manufacturer’s instructions. RNA was then treated with DNase I (Thermo). cDNA was synthesized from a total of 250-1,000 ng of RNA using the NZY Fisrt-Strand cDNA synthesis kit (NZYTech). Primers used for RT-qPCR experiments are listed in Additional File 2D. In all the cases, gene expression was normalized with the housekeeping gene *Act5c* (primers: 5’ –gcgcccttactctttcacca-3’ and 5’ - atgtcacggacgatttcacg-3’. We performed the RT-qPCR analysis with SYBR Green (BioRad) or qPCRBIO SyGreen Blue Mix LO-ROX (PCR BIOSYSTEMS) on an iQ5 Thermal cycler or CFX Real-Time PCR, respectively. Results were analyzed using the dCT method following the recommendations of the MIQE guideline (107).

### Infection survival assays

We performed infection survival assays with mutant, RNAi, overexpression strains comparing their mortality to the mortality of strains with similar genetic backgrounds (Additional File 2A). We also performed infection survival experiments using outbred flies with and without *FBti0019386* and CRISPR/Cas9 mutant flies *FBti0019386-A1* and *FBti0019386-A2*. Female flies were placed in groups of 10 per vial, and we performed the experiments with 5-12 vials (Additional File 2C), except for cn^1^ considered as a wild-type background for which we used 3 vials. Flies were orally infected with *P. entomophila* as explained above. As a control for each experiment, we exposed 3-4 vials containing 10 flies each to sterile LB with 1.25% sucrose. Fly mortality was monitored at several time points until all the flies were dead. Survival curves were analyzed with log-rank test using SPSS v21 software. If the test was significant, we calculated the odds-ratio and its 95% confidence interval when 50% of the susceptible flies were dead, except for *CG8008* and *cbx* that was estimated when 30% and 96% of the susceptible flies were dead.

### Allele-specific expression analysis (ASE)

For each TE analyzed, we first identified two strains homozygous for the presence and two strains homozygous for the absence of the TE according to *T-lex2* or *TIDAL* (50, 51). We then looked for a synonymous SNP linked to the presence of the TE and located in the coding region of the nearby gene. Note that we only selected a SNP when it is present in the coding region of all the alternative transcripts described for that gene. To select the SNP, we downloaded the coding region of the nearby gene from the sequenced DGRP strains available in http://popdrowser.uab.cat/ (108). Once we identified a diagnostic SNP, we re-sequenced the region identified in the used strains to confirm the presence of the SNP, and we performed a PCR to confirm the presence or the absence of the TE. We selected a synonymous SNP that is not linked to the TE in any of the strains analyzed (Additional File 6A). We also analyzed the coding region of the gene in order to discard the presence of nonsynonymous SNPs that could be linked to the TE (Additional File 6B). Additionally, we analyzed the flanking regions of each TE in order to discard other variants that could be linked to the TE, or that could be potentially modifying the gene regulatory regions (Additional File 6C). To do this, we used VISTA to define the conserved regions in the 1 kb TE flanking sequences between *D. melanogaster* and *D. yakuba*, which diverged approximately 11.6 Mya (109). We then checked whether there is any SNP linked to the presence of the TE in the DGRP strains. Only for the *AGO2* gene, we found two SNPs in the coding region that were linked to the TE insertion (Additional File 6B). *AGO2* is a gene showing a fast rate of adaptive amino acid substitutions (74, 110), and it is associated with a recent selective sweep (74). However, it is still not clear which is the genetic variant that is under positive selection (74). Thus, for 13 out of the 14 TEs analyzed, we could not detect any other polymorphism that could be responsible for the observed allele-specific expression differences suggesting that the TE is the most likely causal mutation. We were not able to analyze five of the candidate TEs: for three TEs, *FBti0019381*, *FBti0061105* and *FBti0062242,* we could not identify homozygous strains with and without the TE. For *FBti0019564*, we could not identify a diagnostic SNP. Finally, for *tdn17,* we could not design primers to validate the diagnostic SNP due to the presence of repetitive sequences in the nearby gene.

We then crossed a strain with the TE with a strain without the TE differing by the diagnostic SNP to obtain heterozygous flies in which allele-specific expression was measured (Additional File 6A). Note that for each TE two crosses were performed so that ASE was measured in two different genetic backgrounds. ASE was measured in non-infected and infected conditions. We obtained cDNA samples from three biological replicates. We also extracted genomic DNA (gDNA) from 15-20 heterozygous females for each cross, which is needed to correct for any bias in PCR-amplification between alleles (111). cDNA and gDNA samples were sent to an external company for primer design and pyrosequencing. We analyzed the pyrosequencing results as described in Wittkopp et al (2011) (111). Briefly, we calculated the ratios of the allele with the TE and the allele without the TE of the cDNA samples, and we normalized the values with the gDNA ratio. In order to perform the statistical analysis, we transformed the ratios with log2, and we applied a two-tailed t-test in order to check whether there were allele expression differences between the alleles. We corrected the p-values for multiple testing using Benjamini-Hochberg’s false discovery rate (5% FDR) (112).

### Chromatin immunoprecipitation-qPCR

We performed ChIP-qPCR experiments to detect whether TEs associated with allele specific lower expression were adding H3K9me3 repressive marks (67, 68). For that, we compared the histone mark levels in homozygous flies with the TE with the levels in homozygous flies without the TE. We used y^1^;cn^1^bw^1^sp^1^ strain (113), the strain that was sequenced to obtain the *D. melanogaster* reference genome sequence (114–116), as the homozygous strain with *FBti0020057, FBti0018883*, *FBti0018877, FBti0019985* and *FBti0020137* insertions, and *RAL-908*, as the homozygous strain without those insertions (89). We also checked H3K9me3 marks in *tdn4* using y^1^;cn^1^bw^1^sp^1^ strain as homozygous for the absence and *RAL-810* homozygous for the presence of the insertion. We first confirmed by PCR the presence or absence of each insertion in the strains used. To detect H3K9me3 levels associated to the TE, we designed primer pairs in the TE flanking regions (“left” and “right”): one primer inside the TE sequence and one primer outside the TE sequence (Additional File 7). To detect H3K9me3 levels in the strains without the TE, we used the left forward primer and the right reverse primer. Primer efficiencies ranged from 90-110%. We used a total of 45-55 guts per strain and we performed three biological replicates for each. To obtain the chromatin, we followed Magna-Chip™ A/G kit (from Merck) protocol. After dissection, we homogenized the samples in the buffer A1 with a dounce 30 times, and we crosslinked the guts using formaldehyde at a final concentration of 1.8% for 10 minutes at room temperature. We stopped the crosslink by adding glycine at a final concentration of 125 mM, we incubated samples three minutes at room temperature, and kept them on ice. Then, we washed the samples three times with buffer A1, and we incubated the sample for three hours at 4°C with 0.2 ml of lysis buffer. After lysis, we sonicated the samples using Biorruptor^®^ pico sonication device from Diagenode: 14 cycles of 30 seconds ON, 30 seconds OFF. We kept 20 μl of input chromatin for the analysis (see below), and we immunoprecipitated 80 μl of the remaining sample with antibody against H3K9me3 (#ab8898 from Abcam). As a control for the immunoprecipitation, we checked the H3K9me3 levels in the genes *18S* and *Rpl32* that are expected to be, respectively, enriched and depleted for this histone mark (Additional File 7). We quantified the immunoprecipitation enrichment by RT-qPCR normalizing the data using the “input” of each IP as the reference value using the dCT method as mentioned before.

### In vivo enhancer reporter assays

We generated transgenic flies carrying the TE sequence in front of the *lacZ* reporter gene by using the *placZ.attB* vector (accession number: KC896840) (117). In order to construct a clone with the correct orientation in the promoter region of *lacZ*, two cloning steps were necessary. We first had to introduce specific restriction sites into the flanking regions for each TE sequence. For that, we introduced the restriction sites with the primers used to amplify the region containing the TE sequence (Additional File 8). We used a high fidelity Taq DNA polymerase for DNA amplification (Expand High Fidelity PCR system from Sigma). After that, we cloned the PCR product into the vector pCR4-TOPO^®^ (Invitrogen). Finally, we digested both vectors and ligated the TE sequence into the *placZ.attB*, and we sequenced the cloned insert to ensure that no polymerase errors were introduced in the PCR step. We purified the vector with the GeneEluteTM Plasmid Miniprep kit (Sigma), and prepared the injection mix at 300 ng/μl vector concentration diluted with injection buffer (5 mM KCl, 0.1 mM sodium phosphate, pH 6.8). The injection mix was sent to an external company to inject embryos from a strain that contain a stable integration site (Bloomington stock #24749). After microinjection, surviving flies were crossed in pairs and the offspring was screened for red eye color, which was diagnostic for stable mutants. We established three transgenic strains for each analyzed TE, which were considered as biological replicates in the expression experiments. As a negative control, we also established transgenic strains with the *placZ.attB* empty vector, in order to control for possible *lacZ* expression driven by the vector sequence.

For *FBti0018868*, we designed primers flanking the TE and cloned the PCR product in front of the *lacZ* reporter gene (Additional File 8). For the other two TEs, we constructed two different clones to generate two transgenic strains: one strain with the TE and the other strain without the TE. For the TE *FBti0061506*, which spans only 48 bp, one strain carries the TE and part of the flanking intronic region, and the other strain contains the same genomic region without the TE. For the TE *tdn8,* one strain carries the upstream region of *CG10943*, including the 5’UTR, with *tdn8*, and the other strain carries the same genomic region without *tdn8*. Finally, for the TE *FBti0020057*, we cloned the whole intergenic region, including the UTRs of the flanking genes (Additional File 8).

For the transgenic strains generated in the in vivo enhancer assays, we checked *lacZ* expression in female guts in non-infected and infected conditions. We used the forward primer 5’- cctgctgatgaagcagaacaact-3’, and reverse primer 5’- gctacggcctgtatgtggtg-3’ to check *lacZ* expression. Gene expression was normalized with the housekeeping gene *Act5c*. We performed all RNA extractions, cDNA synthesis and RT-qPCR analysis as mentioned above.

### Immunofluorescence staining

We performed immunofluorescence gut staining to localize β-*GAL* expression in the transgenic flies from the enhancer assays, both in non-infected and infected conditions. Flies were dissected and gut tissue was fixed with 4% Formaldehyde. The tissue was then stained by using the primary antibody mouse anti-βGalactosidase (Hybridoma bank 40-1a), and the secondary antibody anti-mouse Alexa Fluor ® 555 (Sigma). Images were analyzed and captured using a Leica SP5 confocal microscope.

### Transcription factor binding site mutagenesis

Mutagenesis of the binding sites for the predicted *Caudal* and *DEAF-1* transcription factors was performed sequentially with the Q5 Site-Directed Mutagenesis kit (New England Biolabs) taking as a template the *pGreenRabbit* vector containing the *FBti0019386* sequence following manufacturer’s instructions (25). To perform the *Caudal* binding site mutagenesis, we used the primer pair 5’ -actcgatcggacctcact- 3’ and 5’ -gtgtgttagagagagatgacaatg- 3’. To perform the *DEAF-1* binding site mutagenesis, we used the primer pair 5’ -cctctgccgcagcgctcg- 3’ and 5’ -cagcctctgcagctgagtgagg- 3’. Deletions were checked by PCR with the primers 5’- cgacgtgttcactttgcttgt -3’ and 5’- gtaccttcaaatacccttggatcg-3’. PCR bands were confirmed by Sanger sequencing. Vector purification, injection mix preparation, embryo microinjection and fly strain generation were performed as explained above. We then checked *Gfp* expression in female guts in control and infected conditions. We used the forward primer 5’- atgatcagcgagttgcacgcc- 3’, and reverse primer 5’- gacggaaacatcctcggccaca-3’ to check *Gfp* expression. Gene expression was normalized with the housekeeping gene *Act5c*. We performed all RNA extractions, cDNA synthesis and RT-qPCR analysis as mentioned above.

### Detection of alternative transcripts

To confirmed whether *Bin1* transcripts starts in *FBti0019386*, as reported by Batut et al (2013) (24), we performed RT-PCR in the gut tissue of non-infected and infected flies. We used the forward primer 5’- atctgaagctcgttggtggg-3’ and the reverse primer 5’ - atgagactcctgtttcgccg- 3’ to detect *Bin1* transcript starting in the TE, and the same forward primer with the reverse primer 5’- aagagcaaagagaagccggaa- 3’ to detect *Bin1* short transcript.

## DECLARATIONS

### Ethics approval and consent to participate

Not applicable.

### Consent for publication

Not applicable.

### Availability of data and materials

The datasets supporting the conclusions of this article are included within the article and its additional files.

### Authors’ contributions

A.U., M.M., and J.G. designed and managed the project. A.U. and M.M. performed the experiments. A.U., M.M., and J.G. analysed the data and wrote the paper. J.G. revised the paper. All authors read and approved the final manuscript.

## Acknowledgements

We thank members of the González lab for comments on the manuscript.

## SUPPLEMENTARY INFORMATION

**Additional File 1**

**1A**) Reference TE dataset present in genomic regions with recombination rate larger than zero. **1B**) Non-reference TE insertions. **1C**) Candidate adaptive TE dataset. **1D**) Gene functional information for the genes nearby candidate adaptive TEs. **1E)** Immune-related TFBS found in the TEs analyzed in the ASE experiments as reported by Villanueva-Cañas et al (2019).

**Additional File 2**

**2A**) Genotype information on the gene disruption, overexpression, and RNAi stocks used in this work, and summary of the expression and survival assays results. **2B**) Expression results (RT-qPCR) for the gene disruption, overexpression, and RNAi stocks used in this work. **2C**) Survival assay results for the gene disruption, overexpression, and RNAi stocks used in this work. **2D**) Primers used for the RT-qPCR analysis.

**Additional File 3**

Allele ratios from the allele specific expression (ASE) analysis

**Additional File 4**

*D. melanogaster* strains used in the different experiments.

**Additional File 5**

Primers used for the TE screening analysis.

**Additional File 6**

**6A**) Fly strains and SNPs used in the ASE crosses for each gene. **6B**) Analysis of the SNPs in the coding regions of the genes analyzed in the ASE. **6C**) Analysis of the 1 kb TE flanking regions (upstream and downstream) conserved between *D. melanogaster* and *D. yakuba*.

**Additional File 7**

Primers used for ChIP RT-qPCR experiments and H3K9me3 levels in the genes *18S* and *Rpl32*.

**Additional File 8**

Primers used for the amplification of the genomic regions analyzed in the enhancer assays.

